# *De-novo* genome assembly of four rails (Aves: Rallidae): a resource for comparative genomics

**DOI:** 10.1101/2023.12.25.573037

**Authors:** Julien Gaspar, Steve A. Trewick, Gillian C. Gibb

## Abstract

The rails are a phenotypically diverse family of birds that includes around 130 species and displays a wide distribution around the world. Here we present annotated genome assemblies for four rails from Aotearoa New Zealand: two native volant species, pūkeko *Porphyrio melanotus* and mioweka *Gallirallus philippensis*, and two endemic flightless species takahē *Porphyrio hochstetteri* and weka *Gallirallus australis*. The quality checks and comparison with other rallid genomes showed that the new assemblies were of high quality and that the annotations could be trusted. Using the sequence read data, heterozygosity was found to be lowest in the endemic flightless species and this probably reflects their relatively small populations. This study significantly increases the number of available rallid genomes and will enable future genomic studies on the evolution of this family.

## Introduction

Rails (Aves: Rallidae) are a phenotypically diverse family of primarily terrestrial birds with relatively short wings and strong, variably elongated bills (Ripley et al. 1977, Taylor 1998, Livezey 2003). Despite the terrestrial lifestyle of the majority of the species (Taylor 1998), this bird family displays remarkable dispersal capacity resulting in broad distribution and the colonization of numerous oceanic islands (Olson 1973, Ripley et al. 1977, Garcia-R et al. 2017). At the same time, more than 30 flightless rail species are known (Steadman 1995, Kirchman 2012) and a large proportion of them are endemic to single oceanic islands, demonstrating that their ancestors had been volant (Trewick 1997a, b). The large number of flightless species as well as the fact that flightlessness evolved many times amongst extant rails provides a suitable system with which to study genomic changes associated with maintenance and loss of flight in birds.

Rallidae has its origin during the Eocene around 40 million years ago (Garcia–R et al. 2014) and has diversified into over 130 extant species (Steadman 1995, Kirchman 2012, Garcia–R et al. 2014). Rails are part of the order Gruiformes that includes two suborders; the Gruoidea containing, among others, the cranes (family Gruidae) and the Ralloidea that is dominated by the rails (family Rallidae) (Fain et al. 2007, Boast et al. 2019). Phylogenetic analyses of mitochondrial and nuclear genes show that rails comprise eight clades *Fulica, Aramides, Porphyrio, Rallina, Porzana, Laterallus, Gallicrex*, and *Rallus* (Garcia-R et al. 2014).

Despite their phylogenetic diversity (Fig. 1), flightless rails typically exhibit smaller sterna and wings than volant taxa along with wider pelves and more robust femora (Livezey 2003, Gaspar et al. 2020). Moreover, it has been shown that these differences are independent of phylogeny and instead demonstrate convergent evolution associated with a walking ecology (Gaspar et al. 2020). Despite some research using short markers at the population level (Garcia-R. and Trewick 2014, Garcia-R et al. 2017, Trewick et al. 2017), the molecular basis underlying the convergent evolution of flightless rails remain unknown. To investigate that question, more genomic data are needed. Here we present new, high quality, annotated rail genome assemblies of four rail species from Aotearoa New Zealand; two volant, Purple swamphen (called pūkeko in Aotearoa New Zealand) *Porphyrio melanotus* (Temminck, 1820) and buff-banded rail (also called mioweka and moho pererū) *Gallirallus philippensis* (Linnaeus, 1766), and two flightless species, takahē *Porphyrio hochstetteri* (Meyer, 1883) and weka *Gallirallus australis* (Sparrman, 1786). These four genome assemblies were generated to provide two volant-flightless pairs of closely related living species, that will enable future genomic comparisons to highlight the differences and similarities in evolutionary trends between rails with and without the ability to fly.

**Figure 1:**
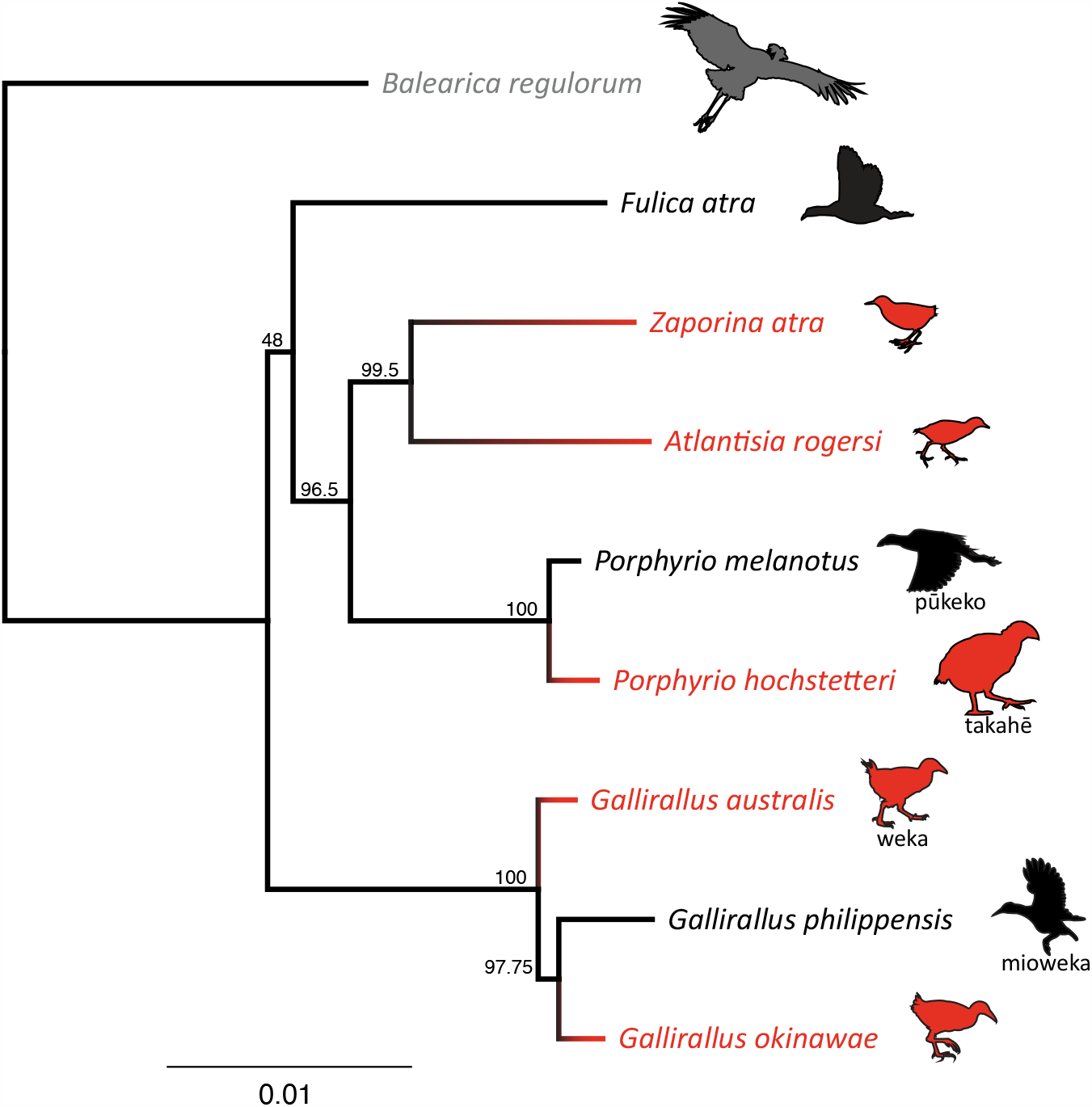
Maximum likelihood (RAxML V.8) phylogeny of three volant (black) and five flightless (red) rail lineages (Aves: Rallidae) based on 10 concatenated nuclear genes analysed with *Balearica regulorum* crane (Aves: Gruidae) (grey) as outgroup; bootstrap supports are indicated for each node.

## Methods

### DNA extraction and sequencing

DNA was extracted from muscle tissue samples of four rails sampled in Aotearoa New Zealand: *Porphyrio melanotus, Gallirallus philippensis, Porphyrio hochstetteri* and *Gallirallus australis*. Extraction used the Geneaid^©^ Genomic DNA Mini Kit following the kit instructions and eluted in 100 μl. DNA quality was then verified by gel electrophoresis and quantified using Qubit 2.0 (Table. 1). Library preparation using the TruSeq Nano DNA kit and quality check were performed by the Massey University Genome Service (New Zealand) with sequencing by Novagene (Hong Kong). Libraries were sequenced on the Illumina HiSeq™ X platform generating non-overlapping 150 bp paired-end reads with an insert size of 550 bp. Fastp V0.19.4 (Chen et al. 2018) was used with default settings for paired-end data to trim the adapters as well as filter and assess the read quality.

**Table 1:** DNA concentration and sampling information including the location and date of collection as well as museum ID (when known)

| Species | DNA concentration | Sampling | Sex | ID |
| --- | --- | --- | --- | --- |
| *Porphyrio melanotus* | 35.9 ng/μl | Roadkill, Turitea Valley near Palmerston North, North Island, New Zealand, within the rohe (area) of Rangitāne o Manawatū. October 2018 | Male | MUNZ12900 |
| *Porphyrio hochstetteri* | 5.0 ng/μl | Provided by the Department of Conservation via Massey University Veterinary Pathology. A translocated individual on Maud Island, Marlborough Sounds, New Zealand | Male | NA |
| *Gallirallus philippensis* | 41.4 ng/μl | Roadkill, Whananāki estuary, Northland, North Island, New Zealand, within the rohe of Ngatiwai. Retrieved March 2011 | Male | MUNZ12901 |
| *Gallirallus australis* | 40.0 ng/μl | Roadkill Granity, West Coast, South Island, New Zealand, within the rohe of Ngāi Tahu. Retrieved July 2012. | Male | MUNZ12767 |

### Genome Assembly

De novo assembly was performed for each of the genomes using Meraculous (Chapman et al. 2011). Average insert size, standard deviation and average read lengths were estimated using sequence reads mapped to a nuclear gene of a close species. Following the Meraculous manual instructions, a range of k-mer sizes was analysed using KmerGenie V1.7051 (Chikhi and Medvedev 2014). The k-mer frequency histograms were reviewed and k for which the main haploid peak had a coverage of at least 30x and a distinct trough to its left that was at most 1/10 of the peak height was chosen. These were 61, 87, 61, 57 for respectively *Porphyrio melanotus, Porphyrio hochstetteri, Gallirallus philippensis* and *Gallirallus australis* (Fig. 2). High heterozygosity for *G. philippensis* meant that completely optimal peak height/trough specs could not be met but the assembly was still successful. See supplementary material for full details of settings used in all Meraculous runs.

**Figure 2.**
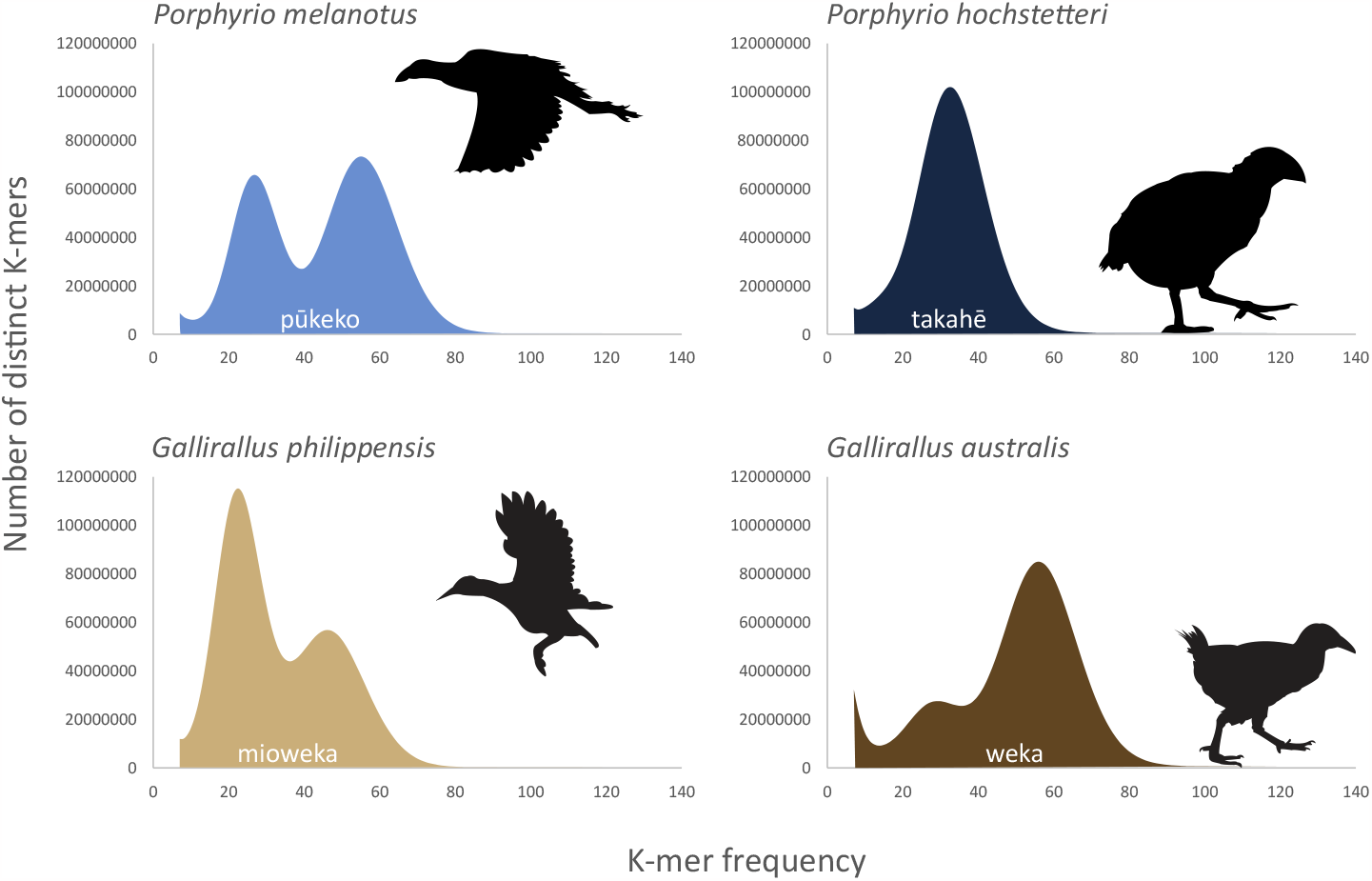
:K-mer frequency in four rails from Aotearoa New Zealand. K-mer (nucleotide sequence of a certain length) were 57, 61, 61, 87 for *G. australis, G. philippensis, P. melanotus* and *P. hochstetteri* respectively. In each distribution, two main peaks correspond to the genomic K-mers for the heterozygous (left) and homozygous (right) parts of the genome. The single main peak of *P. hochstetteri* indicates high homozygosity. Low depth peaks corresponding to erroneous K-mer populations have been masked for clarity. Icons indicate flightless and volant species.

Meraculous (Chapman et al. 2011) was implemented using a docker container we created, which is publicly available at both Github and docker (https://github.com/GenomicsForAotearoaNewZealand/genomics-tools, https://hub.docker.com/r/gfanz/meraculous). The assembly was run through the Catalyst Cloud server (https://catalystcloud.nz) using a cloud instance with 32 vCPU and 256 GB RAM. Runs took between 1 and 3 days per assembly.

### Additional Genomes

In order to assess the quality of our genome assemblies, we compared them to a selection of additional rail genomes, Okinawa rail *Gallirallus okinawae (also known as Hypotaenidia okinawae)*, GenBank assembly accession: GCA_027925045.1, takahē *Porphyrio hochstetteri* GCA_020800305.1, Henderson crake *Zapornia atra* (formerly *Porzana atra*) GCA_013400835.1, Eurasian coot *Fulica atra* GCA_013372525.1, and Inaccessible Island rail *Atlantisia rogersi* GCA_013401215.1 The genome of a grey crowned crane *Balearica regulorum* (order Gruiformes, family Gruidae; Bennett, 1834) GCA_011004875.1 was used as a reference for the gene annotations.

### Quality assessment

Meraculous outputs were used to compare the sequence length of the shortest scaffold at 50% of the total genome length (N50) and the smallest number of scaffolds whose total length makes up half of the genome size (L50) values as well as the assembly length and the number of contigs and scaffolds. Busco v4 (Seppey et al. 2019) was implemented using a Docker (Merkel 2014) container (default parameters, mode: genome) on the genomes using the aves_odb10 dataset to assess the assembly completeness.

### Genome annotation

Geneious R.11 (https://www.geneious.com) was used to extract the coding sequences (CDS) from *B. regulorum* genome (GCA_000709895) and these were filtered to retain only the longest CDS per gene where multiple annotations existed. Gmap (version 2019-09-12) (Wu and Watanabe 2005) was used to annotate the newly assembled genomes. Each assembly was first indexed using the gmap_buil function, then *B. regulorum* CDS were mapped to it with the setting -f 2 to obtain a GFF3 formatted annotation.

### Extracting coding regions

During the assembly process, exons from the same gene are sometimes assembled into different scaffolds. To obtain a sequence list containing the entire coding region for each gene, the exons were extracted using Geneious R.11 and remapped to the *B. regulorum* CDS with BWA (0.7.17-r1188) using BWA-mem with the default settings (Li 2013).

To assess the size and quality of the extracted CDS for each genome they were compared to the *B. regulorum* reference. The quality (complete or partial) of coding regions retrieved was assessed using the samtools V.1.9 (Li et al. 2009) faidx tool (to obtain the length of each sequence) and a custom R script to compare the CDS sequences with the reference (see supplementary data).

### Heterozygosity

Read depth, coverage and heterozygosity of the newly assembled genomes were estimated using twenty randomly selected genes (*ADA, DHX40, ENPEP, EXOG, FAM196B, FUBP3, GOLGA7B, GRHL3, KCNK5, LEMD3, LOC104630315, LOC104633950, LOC104643156, MLNR, MMS19, PIANP, THOC3, ZCCHC2, ZNF410*, and *ZRANB1*) for a total length of 266,456 bp and the paired reads for each genome mapped to them in Geneious R.11 with low sensitivity/fast mapping settings. The Geneious ‘Find variations/SNPs’ tool in the ‘Annotate & Predict’ section was used with the following settings: minimum coverage of 50 and minimum variant frequency of 0.3 to locate the heterozygous sites. Heterozygosity was then estimated by dividing the number of heterozygous sites by the total length of the concatenated gene sequences. This method, despite not using the whole genome to assess the heterozygosity level of each species, generates reliable estimates that can be compared between lineages.

### Phylogeny

Phylogenetic inference to show relative relationships between the four new genomes and other selected rails with *Balearica regulorm* as outgroup was performed using 10 genes selected from a set of universal nuclear markers suitable for avian phylogenetic reconstruction (Liu et al. 2018). The genes were *ADNP, BEGAIN, INO80D, KBTBD8, NCOA6, RHOBTB1, S1PR3, SPECC1L, ZNF618* and *ZNF654*. These 10 CDS alignments were concatenated into a 21,390 bp alignment using Phyluce v1.7.1 (Faircloth 2016) with the default settings and the best-fit partitioning scheme was determined using PartitionFinder2 (Lanfear et al. 2017) via the CIPRES Science Gateway (Miller et al. 2010). A list of genes and partitions can be found in the supplementary data). Maximum Likelihood (ML) analyses were implemented in RaxML v8.2.10 (Stamatakis 2014) via the CIPRES Science Gateway with bootstrapping automatically stopped employing the majority rule criterion. The consensus tree was then visualized in Geneious (Fig. 1).

## Results

### DNA extraction and sequencing

The raw data comprised between 780 million (*G. philippensis*) and 936 million (*G. australis*) paired reads per species. Most of these were retained after the filtering and cleaning step (Table 2). Fastp generates a Phred quality score (Q score) for each of the species that represents the ratio of bases with a probability of containing no more than 1/100 (Q20) or in 1/1000 (Q30) errors (Ewing and Green 1998, Ewing et al. 1998, Richterich 1998). These scores range between 97.37% and 98.5% for Q20 and between 93.74% and 95.23% for Q30 implying high sequencing quality for all four species.

**Table 2:** Fastp outputs after the sequencing of four rail species indicating the number of reads before and after filtering as well as the quality assessment.

| Species | Before fastp filtering | After fastp filtering |  |  |  |
| --- | --- | --- | --- | --- | --- |
|  | Total reads | Total reads | % reads conserved | Q20 bases | Q30 bases |
| *P. melanotus* | 894.570034 M | 881.624382 M | 98.55% | 97.73% | 94.46% |
| *P. hochstetteri* | 845.999032 M | 817.395666 M | 96.62% | 97.37% | 93.84% |
| *G. philippensis* | 781.084610 M | 760.059606 M | 97.31% | 97.39% | 93.74% |
| *G. australis* | 936.861886 M | 917.214824 M | 97.90% | 98.05% | 95.23% |

K-mer frequency plots can be used to estimate the level of heterozygosity for each individual and by proxy each species. Indeed, k-mers from the heterozygous regions (left peak on Fig. 2) will have half the sequencing coverage (i.e., K-mer frequency) compared to the homozygous regions (right peak). The higher the left peak the higher the heterozygosity. The two volant species *G. philippensis* and *P. melanotus* exhibited high heterozygosity with the left peak being higher than the right for *G. philippensis*. A very low left peak was found for the *G. australis* data and only one peak was observed for *P. hochstetteri*. This implies a much lower level of heterozygosity for both of the endemic, flightless species that have limited populations.

To compare heterozygosity between the newly assembled genomes, paired reads were mapped to a set of 20 genes for each species and the ratio of heterozygous sites divided by the total sequence length was calculated (Fig. 3). The two volant species showed a higher heterozygosity level than the two flightless species. Based on the paired reads mapping, the mean depth of coverage was calculated for each species (Table. 3) with the overall average being 96.4x.

**Figure 3:**
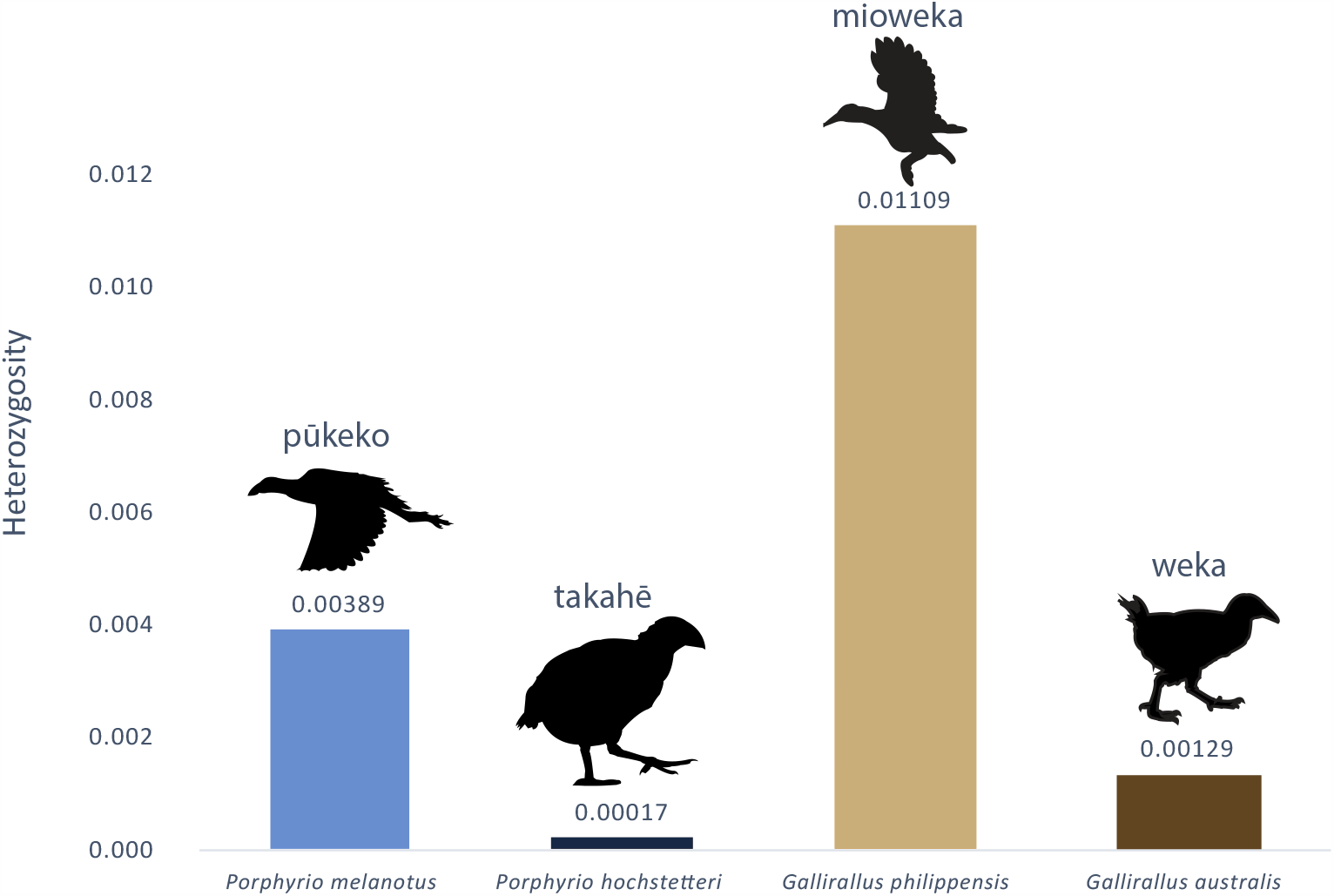
Average heterozygosity at 20 randomly selected genes from four newly assembled and annotated rail genomes (average total length 266,456 bp). Heterozygosity is the proportion of total nucleotide sites per individual site having two bases. Icons indicate flightless and volant species.

### Genome assembly

Meraculous de novo assemblies yielded scaffold N50 between 126 kb (*G. australis*) and 30 kb (*P. hochstetteri*) and scaffold L50 between 2,365 (*G. australis*) and 6,047 (*G. philippensis*) (Table.2). The total genome assembly size of the four newly assembled rails differed little with a range from 1.07 Gb (*G. philippensis*) to 1.16 Gb (*G. australis*). This was similar to the previously assembled rails (between 1.11 and 1.27 Gb, see Table 2) and slightly shorter than the crane *B. regulorum* (1.22 Gb).

Busco scores were similar for *P. melanotus, P. hochstetteri* and *G. australis* with close to 80% of single copy genes were found complete. In contrast, *G. philippensis* comprised 69% of “Complete single copy” and had a higher proportion (17%) of missing genes (Fig. 4).

**Figure 4:**
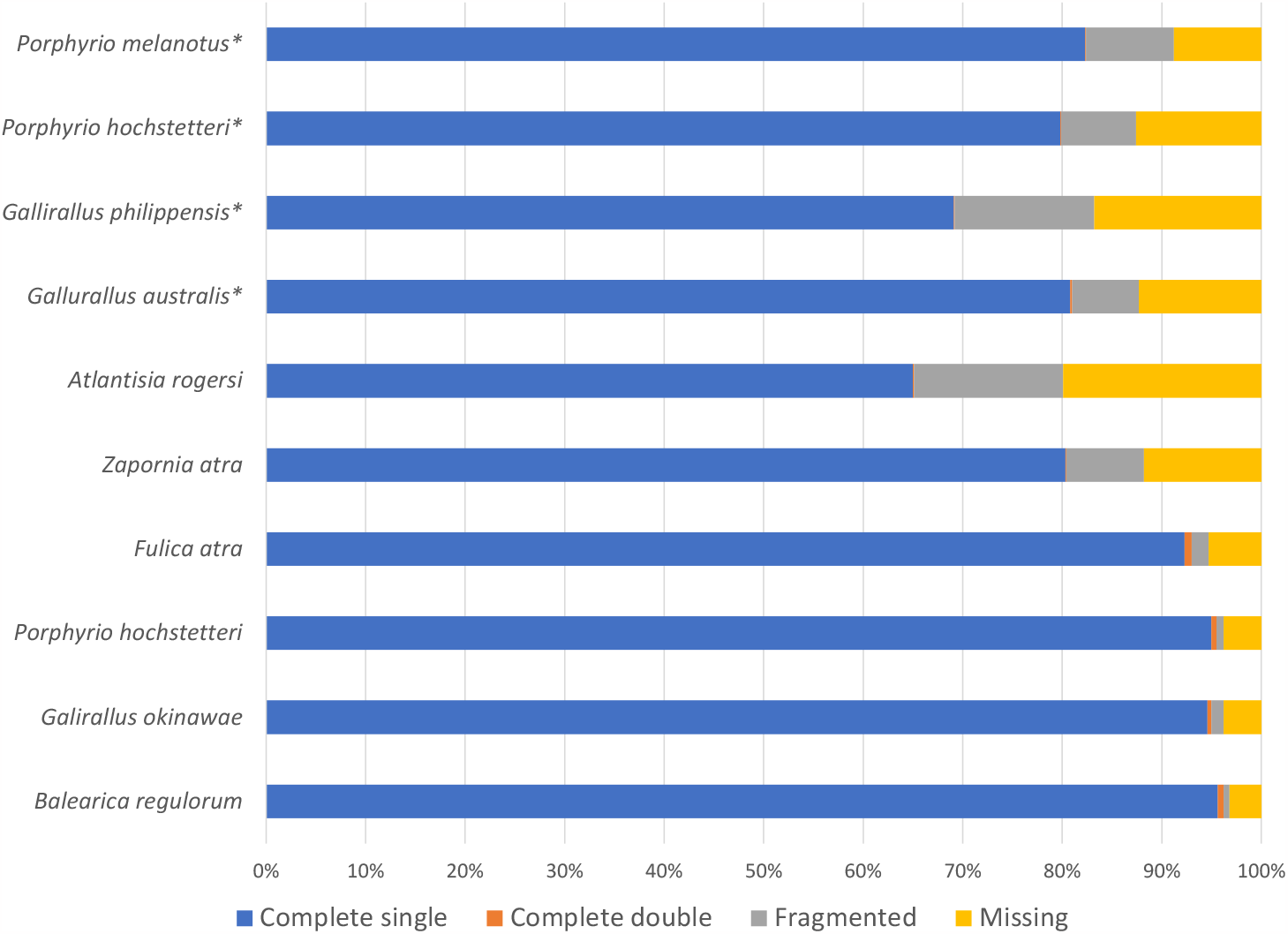
BUSCO V4 results (mode: genome) using the aves_odb10 dataset. Total number of genes (Benchmarking Universal Single-Copy Orthologs): 8338. Complete: ortholog is present in the genome in a single copy (Complete single) or in two copies (Complete double), Fragmented: gene is only partially present, Missing: no significant match in the genome. *New assemblies

### Extracting coding regions

For the four new rail assemblies, the coding regions of each gene were extracted based on the annotations and compared with the respective *B. regulorum* CDS. Over 9000 gene CDSs were retrieved near-complete (above 95% of the reference CDS nucleotide sequence length) for the two *Porphyrio* species and *Gallirallus australis* (Fig. 5). *G. philippensis* exhibited a slightly lower proportion (8,259 CDS over 95%) which was consistent with the BUSCO results. The CDSs present in the reference genome but not in the rail data (“Not found” in Fig. 5) represent less than 7.5% of the CDS for all species.

**Figure 5:**
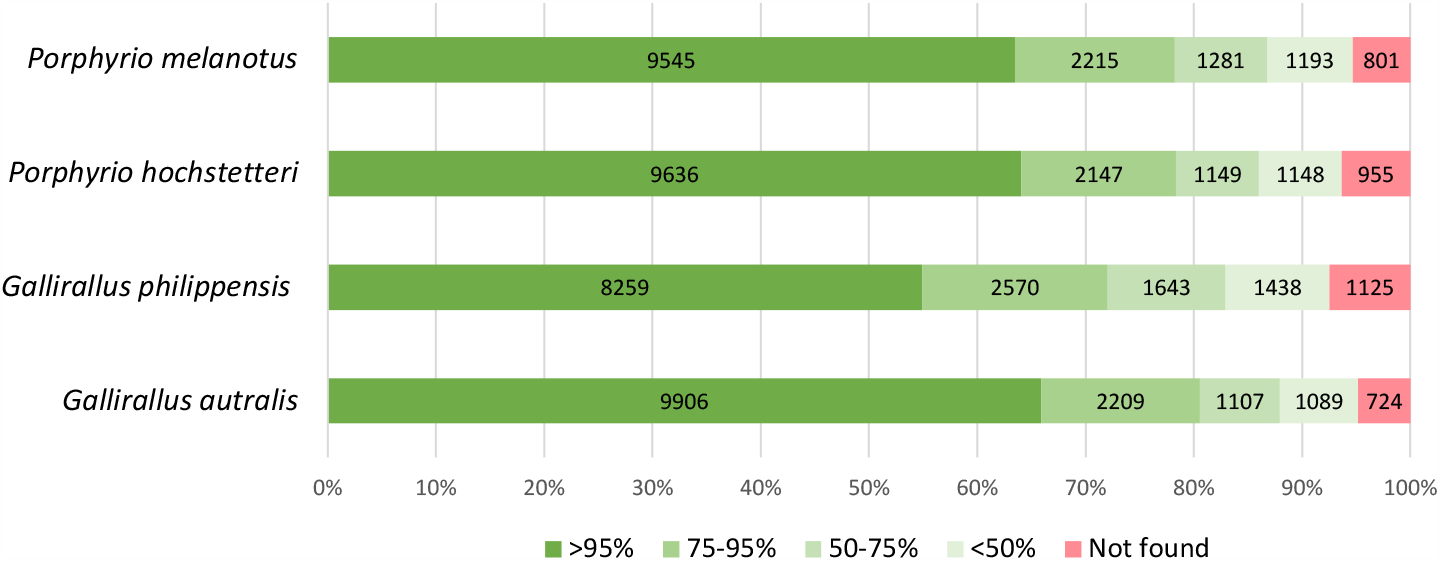
Completeness of CDSs retrieved from eight rail genomes compared to the reference crane *B. regulorum* genome that has a total of 15,035 annotated CDSs. Colours indicate the proportion of genes retrieved from a sample at various scales of completeness.

## Discussion

Considerable variation was observed between species heterozygosity (Fig. 2 and 3). Indeed, the two volant species were more heterozygous than the flightless ones (Fig. 3) with big differences observed between the most heterozygous species, *G. philippensis* (frequency of heterozygous site of 0.01) and the least heterozygous species *P. hochstetteri* (0.0002). The low level of heterozygosity in flightless species probably reflects the relative isolation and reduced size of the habitat as well as population collapse (Baker et al. 1995, Burga et al. 2017, White et al. 2018). The takahē *P. hoschetteri* is a critically endangered flightless species with a population of only 500 in 2023 (www.doc.govt.nz), all derived from a remnant discovered in the 1950s that may have numbered as low as two individuals (Wallace 2002). The resulting inbreeding depression likely explains its extremely low level of heterozygosity (Grueber et al. 2010). *Gallirallus philippensis* on the other hand is a relatively abundant species with a geographic range that covers the islands of Aotearoa New Zealand and the western Pacific (Trewick 1997b, Garcia-R et al. 2017) which is likely to maintain high heterozygosity at the species level.

The four newly assembled genomes have similar or better characteristics than the other rail genomes assembled from Illumina HiSeq data (Fig. 4 and 5, Table 3) with N50 and L50 scaffolds within the same range as these other rails. The BUSCO results (Fig. 4) and CDS extractions (Fig. 5) showed similar trends and add to our confidence that the genome assemblies are of good quality with limited assembly errors. Despite being naturally more fragmented than those assembled using long-read sequencing technology (Table 3), a significant proportion of full-length coding regions were identified and extracted showing good utility for future comparative analyses (Fig. 5).

**Table 3:** De novo genome assembly metrics among 8 rail species and one crane (*B. regulorum*). *New assemblies.

| Species | Genome Assembly size (Gb) | Largest scaffold | Number of scaffolds | Scaffold N50 | Scaffold L50 | Number of contigs | Contig N50 (Kb) | Contig L50 | Depth of coverage | # Ns in assembly (Kb) | Sequencing technology |
| --- | --- | --- | --- | --- | --- | --- | --- | --- | --- | --- | --- |
| *Porphyrio melanotus* * | 1.114 | 1,068,914 | 34,563 | 82,204 | 3,707 | 159,218 | 16.5 | 16,605 | 102.26 | 24,745 | Illumina HiSeq |
| *Porphyrio hochstetteri* * | 1.116 | 1,015,032 | 30,278 | 30,278 | 3,641 | 76,213 | 40.8 | 7,446 | 94.53 | 8,974 | Illumina HiSeq |
| *Gallirallus philippensis* * | 1.07 | 722,688 | 55,205 | 46,737 | 6,047 | 209,712 | 9.6 | 28,118 | 103.04 | 26,818 | Illumina HiSeq |
| *Gallirallus australis* * | 1.158 | 1,692,012 | 36,524 | 126,032 | 2,365 | 96,978 | 41.2 | 7,224 | 85.59 | 13,634 | Illumina HiSeq |
| *Atlantisia rogersi* | 1,168 | 1,015,111 | 159,311 | 36,139 | 8,295 | 160,845 | 36,1 | 8,295 | 41 | 106 | Illumina HiSeq |
| *Zapornia atra* | 1.119 | 1,795,565 | 58,849 | 134,191 | 2,049 | 113,655 | 44,130 | 6,609 | 45 | 10,181 | Illumina HiSeq |
| *Fulica atra* | 1,167 | 27,139,163 | 17,827 | 6,390,841 | 46 | 31,348 | 246,1 | 1,314 | 53 | 19,451 | Illumina NovaSeq |
| *Porphyrio hochstetteri* | 1.27 | 224,114,340 | 173 | 71.6 MB | 5 | 500 | 13.5 MB | 31 | 36.8 | 5,316 | PacBio Sequel II HiFi; Bionano Genomics DLS; Illumina HiSeq; Arima Genomics Hi-C v2 |
| *Galirallus okinawae* | 1.179 | 218,223,205 | 258 | 101.8 MB | 4 | 440 | 20,700 | 16 | 100 | 18 | Illumina Novaseq6000; ONT PromethION |
| *Balearica regulorum* | 1.221 | 219,267,915 | 104 | 82,577,926 | 5 | 248 | 23.3 | 14 | 59.6 | 4,796 | PacBio Sequel I CLR; Illumina NovaSeq; Arima Genomics Hi-C; Bionano Genomics DLS |

Among the four newly assembled rallid genomes, *G. philippensis* had the lowest proportion of complete genes according to both the BUSCO (Fig. 4) and extracted CDS comparison (Fig. 5). This can be attributed to the high heterozygosity level which generally makes the assembly process more challenging due to the increased complexity of the de Bruijn graph structure (Kajitani et al. 2014). Nonetheless, the *G. philippensis* genome is a high-quality assembly that can be used to investigate evolutionary processes along with the three other assembled genomes. For all four genomes, the annotation process was stable, and a large majority of the genes were retrieved. In all species, over 70% of the genes were identified with greater than 75% completeness.

To conclude, we provide here four high-quality assemblies which represent valuable genomic resources to investigate evolutionary processes within the rail family. The quality checks that were performed showed that the generated assemblies are reliable and that the annotations can be trusted. Comparing the newly assembled genomes showed lower levels of heterozygosity in flightless species which likely reflects their relatively small populations. This study significantly increases the number of available rallid genomes, targeting flying-flightless pairs; this creates new opportunities to investigate the evolution of avian flightlessness.

## Data availability

The genomes and annotations are available on NCBI, BioProject PRJNA782688. The configuration files, command lines used, CDS lists, and R scripts are available in the supplementary data. URL: https://figshare.com/s/3a89eea20c4607abbefe.

## Acknowledgements

This study was supported by the New Zealand Marsden Fund Council from Government funding, managed by Royal Society Te Apārangi, grant MAU1601 to GCG. Thanks to Roger Moraga for initial bioinformatic discussions and assistance using the software Meraculous. The genome assemblies were generated with the help of Genomics for Aotearoa New Zealand (GFANZ, genomics.nz) thanks to Rob Elshire. The authors would like to thank Richard Witehira who provided the helpful local knowledge about the mioweka name. Thanks also to Jonathan Proctor (Rangitāne o Manawatū) for ongoing consultation around the role of Rangitāne o Manawatū as kaitiaki (guardians) of the museum collection samples held by Massey University Palmerston North.

